# Elevated Expression of a Functional Suf Pathway in the *E.coli* BL21(DE3) Cell Line Enhances Recombinant Production of an Iron-Sulfur Cluster Containing Protein

**DOI:** 10.1101/811059

**Authors:** Elliot I. Corless, Erin L. Mettert, Patricia J. Kiley, Edwin Antony

**Affiliations:** Department of Biological Sciences, Marquette University, Milwaukee, Wisconsin 53201, USA; Department of Biomolecular Chemistry, School of Medicine and Public Health, University of Wisconsin-Madison, Madison, Wisconsin, 53706, USA; Department of Biochemistry and Molecular Biology. Saint Louis University School of Medicine, St. Louis, MO 63104, USA

**Keywords:** Fe-S protein overexpression, E. coli B, iron sulfur biogenesis, Suf

## Abstract

Structural and spectroscopic analysis of iron-sulfur [Fe-S] cluster-containing proteins is often limited by the occupancy and yield of recombinantly produced proteins. Here we report that *Escherichia coli* BL21(DE3), a strain routinely used to overexpress [Fe-S] cluster-containing proteins, has a nonfunctional Suf pathway, one of two *E. coli* [Fe-S] cluster biogenesis pathways. We confirmed that BL21(DE3) and commercially available derivatives carry a deletion that results in an inframe fusion of *sufA* and *sufB* genes within the *sufABCDSE* operon. We show that this fusion protein accumulates in cells but is - inactive in [Fe-S] biogenesis. Restoration of an intact Suf pathway combined with enhanced suf operon expression led to a remarkable (~3-fold) increase in the production of the [4Fe-4S] cluster-containing BchL protein, a key component of the dark-operative protochlorophyllide oxido-reductase complex. These results show that this engineered ‘SufFeScient’ derivative of BL21(DE3) is suitable for enhanced large-scale synthesis of an [Fe-S] cluster-containing protein.

**IMPORTANCE:** Large quantities of recombinantly overexpressed iron-sulfur cluster-containing proteins are necessary for their in-depth biochemical characterization. Commercially available *E. coli* strain BL21(DE3) and its derivatives have a mutation that inactivates the function of one of the two native pathways (Suf pathway) responsible for cluster biogenesis. Correction of the mutation, combined with sequence changes that increase Suf expression can increase yield and cluster occupancy of [Fe-S] cluster-containing enzymes, facilitating the biochemical analysis of this fascinating group of proteins.

## INTRODUCTION

Iron-sulfur [Fe-S] proteins are integral to the activity of numerous biological processes including respiration, nitrogen fixation, photosynthesis, DNA replication and repair, RNA modification, and gene regulation(1, 6, 10). In *Escherichia coli* K-12, there are two multiprotein systems, Isc and Suf, dedicated to the biosynthesis of various [Fe-S] clusters and their incorporation into 140 known iron-sulfur enzymes, stressing the importance of optimized protocols for their incorporation. (2, 3, 26, 29). The Isc system is encoded by the *isc* operon, composed of the *iscRSUA*-*hscBA*-*fdx*-*iscX* genes (Fig. 1a). The Suf system is encoded by its cognate *sufABCDSE* (*suf*) operon (Fig. 1a). *E. coli* carrying defects in both systems are not viable due to a non-functional isoprenoid biosynthetic pathway which relies on two [Fe-S] enzymes, highlighting the significance of these [Fe-S] cluster biogenesis systems for essential life processes(32). However, the Isc and Suf systems display functional redundancy, as cells lacking only one system remain viable. Nevertheless, individual enzyme components of the two systems are not interchangeable, reinforcing that the scaffolds for building [Fe-S] clusters are functionally different(27, 33). Under normal growth conditions, the Isc system is thought to play the major role in [Fe-S] cluster biogenesis, but under conditions of stress, such as oxidative stress or iron-limiting conditions, the Suf system is reported to assume a greater role(15). Interestingly, some bacteria, archaea, and plant plastids contain only the Suf machinery, serving as the sole [Fe-S] cluster biogenesis machinery(2, 3, 26, 29, 30).

**Figure 1.**
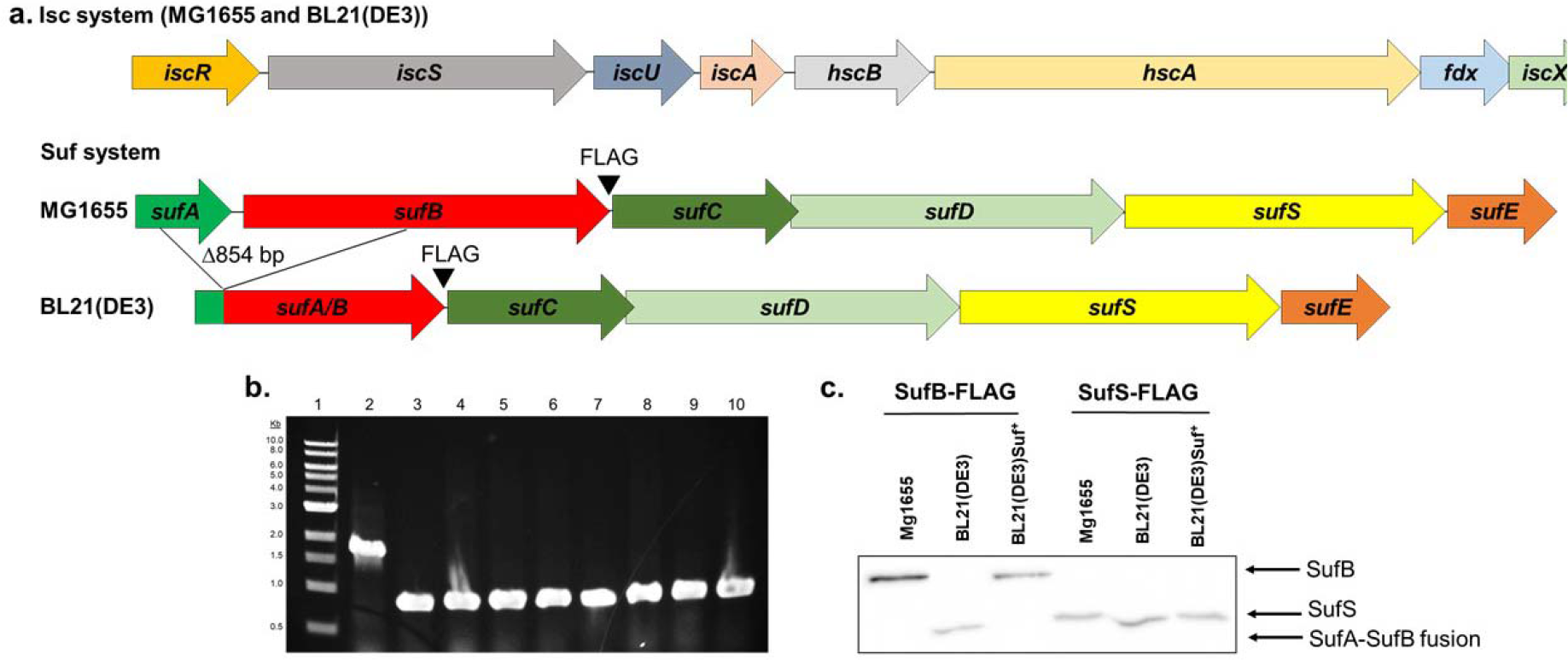
An inframe deletion between *sufA* and *sufB* render the *suf* operon inactive in BL21(DE3). a. Diagram of the *isc* and *suf* operons present in MG1655 and BL21(DE3); BL21(DE3) has a 854 bp deletion in the *suf* operon resulting in an inframe fusion of the *sufA* and *sufB* genes. b. The presence of the 854 bp deletion was tested in commercial lineages of BL21(DE3) using PCR analysis. Lane 2 shows the expected 1641 bp product from MG1655. Lanes 3 through 10 show the 787 bp product predicted from the 854 bp deletion present in strains BL21(DE3), NiCo21(DE3), Lemo21(DE3), C41(DE3), Rosetta2(DE3)pLysS, BLR(DE3)pLysS, BL21(DE3)Ai, and BL21(DE3)codon plus. c. Western blot analysis using an anti-FLAG antibody reveals the production of full length SufB protein in MG1655 and in strain BL21(DE3)Suf^+^, in which the *sufA sufB* genes are properly restored. Full length SufS protein is present in all strains.

To accelerate biochemical studies of [Fe-S] proteins, genes encoding proteins of interest are often heterologously expressed in engineered *E. coli* strains designed for overproduction of proteins. A major challenge in the field is to obtain large enough quantities of proteins at high concentrations that are also maximally occupied with [Fe-S] clusters(34). Increasing the expression levels of housekeeping Isc pathway imparts variable improvement in [Fe-S] cluster protein yields(8, 13, 20, 34). However, to our knowledge, a similar approach has not been examined for the Suf pathway despite it being the sole pathway for [Fe-S] biogenesis in many organisms.

A commonly used strain for [Fe-S] protein overexpression is *E. coli* BL21(DE3) or its derivatives. The ancestry of the parent strain for the modern day BL21(DE3) can be traced back to *E. coli* B strains derived from Delbrück and Luria dating back to the 1920s(5). The sequence of the BL21(DE3) genome, published in 2009, revealed many sequence changes compared to another *E. coli* B strain, REL606(5, 9, 31). Amongst these differences was an in-frame deletion between *sufA* and *sufB* within the *suf* operon, encoding the Suf [Fe-S] biogenesis pathway. Here we show that BL21(DE3) is defective for Suf dependent [Fe-S] biogenesis and we corrected the deletion with sequences found in *E. coli* K12. We also tested commercially available BL21 derivatives to determine if they carry the same deletion of *sufAB*, which was suggested to arise from UV treatment early in the lineage of BL21(31). By altering the promoter sequences of the corrected allele in BL21(DE3) we developed a strain with increased expression of the Suf pathway and tested whether it improved the yield of [Fe-S] proteins upon overexpression. This strain may be of general use in applications that require overexpression of [Fe-S] proteins for structural and spectroscopic studies where large quantities of protein are required.

## RESULTS

### Confirming the in-frame partial deletion of *sufA* and *sufB* in BL21(DE3) and in commercial derivatives

Previous sequencing of the BL21(DE3) genome indicated that it contains a genomic deletion encompassing portions of *sufA*, which functions in delivering [Fe-S] clusters to apoproteins, and *sufB*, which acts as a scaffold for building [Fe-S] clusters(31). PCR amplification of the *suf* operon from our laboratory stock of BL21(DE3) using primers flanking the two genes generated a DNA fragment that is 854 bp smaller than the size expected for intact *sufA* - *sufB* observed for the reference strain *E. coli* K-12 MG1655 (Fig. 1b) indicating the presence of the deletion. We also tested whether other commercially available strain derivatives of BL21(DE3) have a similar deletion. PCR amplification of the *sufA* - *sufB* genes from 7 different commercial strains [Ni-Co21(DE3), Lemo21(DE3), C41(DE3), Rosetta2(DE3)pLysS, BLR(DE3)pLysS, BL21(DE3)Ai, and BL21(DE3)codon plus] revealed the same 854 bp deletion in the *sufA* - *sufB* genes as observed for the parent BL21(DE3) (Fig. 1b).

### The 854bp deletion within the *suf* operon renders the Suf pathway nonfunctional

DNA sequencing of *sufA* and *sufB* of BL21(DE3) also confirmed the same nucleotide deletion boundaries within *sufA* and *sufB* as reported previously. Comparison of the *sufABCDSE* operon DNA sequence from *E. coli* K12 strain MG1655 and *E. coli* B strain BL21(DE3) indicated that the 854bp inframe deletion within the BL21(DE3) *sufABCDSE* operon encompassed the last 79 codons of *sufA* and the first 202 codons of the *sufB* coding sequences, generating a predicted SufA/B fusion protein of ~37 kDa (Fig. 1a). A FLAG epitope was engineered at the C-terminal end of SufB to test whether this fusion protein accumulated in cells by Western blotting using an anti-FLAG antibody. In the BL21(DE3) strain background, a protein that was smaller than that present in the reference strain MG1655 was detected, consistent with the size of the predicted fusion protein between SufA and SufB (Fig. 1c). This deletion had no effect on downstream SufS expression since similar levels of FLAG-tagged SufS were observed comparing MG1655 and BL21(DE3) (Fig. 1c). Activity of this mutant fusion protein was tested using P1 vir to transduce into BL21(DE3) a Δ*iscSUAhscBAfdx*∷*kan* allele which requires a functional Suf pathway for growth(14). No BL21(DE3) derivatives were recovered, suggesting that this fusion protein, and consequently, the Suf pathway, is non-functional in this strain. A derivative of BL21(DE3) was then constructed in which the mutant *suf* operon was replaced with an intact *suf* operon from MG1655 using P1 vir transduction. Instead of the fused protein observed in BL21(DE3), this genetically restored BL21(DE3)Suf^+^ strain (PK13235) generates full length SufB as confirmed by Western blot analysis (Fig. 1c).

### Designing a functional Suf-containing strain for [Fe-S] protein overexpression

Since we and other laboratories routinely utilize BL21(DE3) to overexpress recombinant [Fe-S] cluster-containing proteins, we tested whether restoring the Suf system would improve the yield of recombinant [Fe-S] cluster-containing proteins. Because the *suf* operon is repressed by the transcriptional regulator Fur under standard growth conditions (26, 27), we constructed a variant of BL21(DE3)Suf^+^ that also contains a mutated Fur binding site (designated here as Suf^++^) within the *sufA* promoter region. This mutation increases *suf* expression at least 4-fold under aerobic conditions(14). Thus, we expect this BL21(DE3)Suf^++^ strain (PK11466) to have enhanced levels of the Suf machinery, in addition to functional SufA and SufB proteins. In contrast to BL21(DE3), viable colonies of BL21(DE3)Suf^+^ and BL21(DE3)Suf^++^ were recovered upon deletion of the *isc* operon, indicating that the Suf pathway is indeed functional in these two strains.

### [Fe-S] protein yields, occupancy, and growth times of BchL-overexpressing cultures are improved with restored and upregulated Suf pathway

Expression and yield of recombinant [Fe-S] cluster-containing proteins was assessed in BL21(DE3)Suf^+^ and BL21(DE3)Suf^++^ to determine if the restoration and/or up-regulation of the *suf* operon had any beneficial effects on [Fe-S] cluster protein synthesis and/or incorporation. We chose two protein complexes from the Dark-operative Protochlorophyllide Oxido-Reductase (DPOR) enzyme from *Rhodobacter sphaeroides*, which has only the Suf system for [Fe-S] biogenesis, as a test system. DPOR catalyzes the ATP-dependent reduction of protochlorophyllide (Pchlide) to chlorophyllide (Chlide) in plant and photosynthetic bacterial systems under low-light or dark conditions(22, 25). DPOR is comprised of two components (Fig. 2a): an electron donor (BchL) and an electron acceptor enzyme (BchN-BchB)(4, 24). BchL exists as a homodimer stabilized by a bridging [4Fe-4S] cluster ligated by 2 cysteine residues from each monomer(4, 19). BchN and BchB exist as an A_2_B_2_ hetero-tetramer with two symmetric halves (Fig. 2a). Each half contains one [4Fe-4S] cluster that accepts an electron from reduced BchL and an active site for substrate (Pchlide) binding and reduction(18). Two rounds of electron transfer from BchL to BchN-BchB are required to reduce the C17=C18 double bond in Pchlide to form Chlide(23).

We compared BchL and BchN-BchB synthesis, protein yield, iron content, and protein activity from three strains: a) BL21(DE3), b) BL21(DE3)Suf^+^, and c) BL21(DE3)Suf^++^. Strains containing plasmids carrying open reading frames for BchL or BchN-BchB under the control of an IPTG-inducible T7-promoter were grown and used for synthesis and purification of the proteins under identical conditions. For BchL, overall protein synthesis was similar between BL21(DE3) and BL21(DE3)Suf^+^(Fig. 2b, c) but markedly higher for BL21(DE3)Suf^++^ (Fig. 2d). Overexpressed BchL protein was then purified using affinity Ni^2+^-NTA chromatography. The enhanced overexpression resulted in a reproducible ~3-fold increase in the overall yield of purified BchL protein obtained from one liter of BL21(DE3)Suf^++^ (3.62±0.54 mg/L) compared to BL21(DE3) (1.32±0.63 mg/L) or BL21(DE3)Suf^+^ cells (0.81±0.23 mg/L) (Fig. 3a). We believe this increased yield is due to increased stability of BchL with loaded intact, fully formed clusters. In cell lines lacking the restored Suf pathway, there may be more improperly folded BchL due to decreased overall cluster availability, maintenance, chaperone activity, overall cell health, or a combination of these factors.

**Figure 2.**
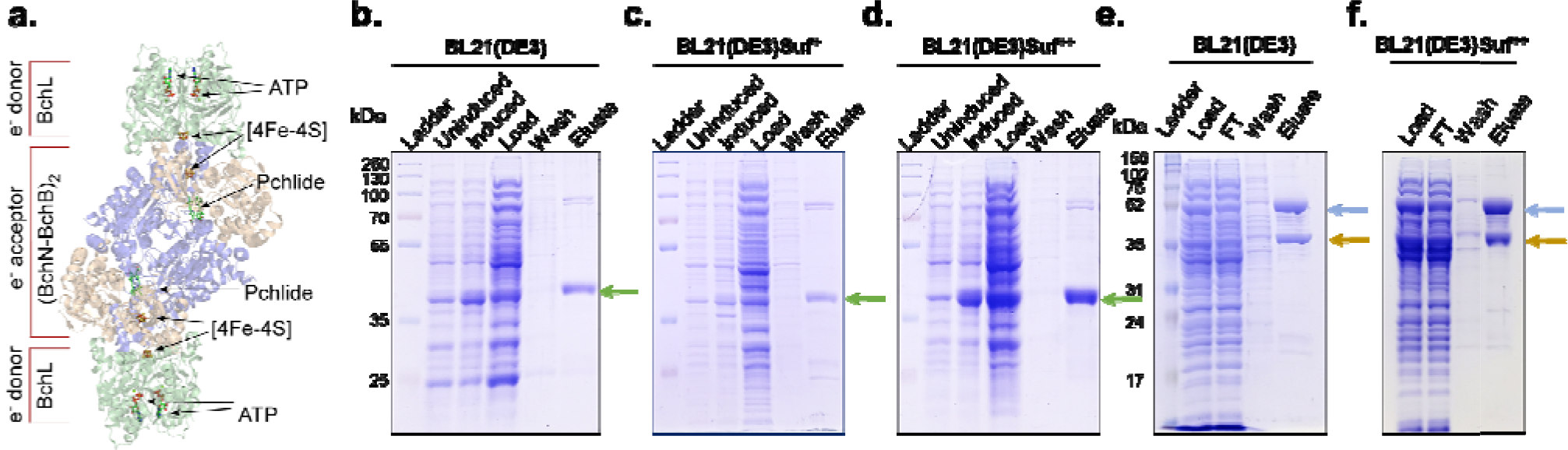
Removing transcriptional repression of the *suf* operon results in enhanced production of the [4Fe-4S] cluster carrying BchL protein in BL21(DE3) cell lines. a) Crystal structure of the DPOR complex (PDB ID:2YNM) consisting of BchL (green) and BchB(blue)-BchN(brown) proteins. Locations of [4Fe-4S] clusters and the binding sites for Pchlide and ATP are denoted. Overexpression and affinity purification of BchL from BL21(DE3), c) BL21(DE3)Suf^+^, and d) BL21(DE3)Suf^++^ cells. Green arrow indicates the position of BchL following SDS-PAGE. Analogous purification of the BchN-BchB complex from e) BL21(DE3), f) and BL21(DE3)Suf^++^ cells. Brown and green arrows indicate position of BchN and BchB respectively. Gels are representative of three independent experiments.

**Figure 3.**
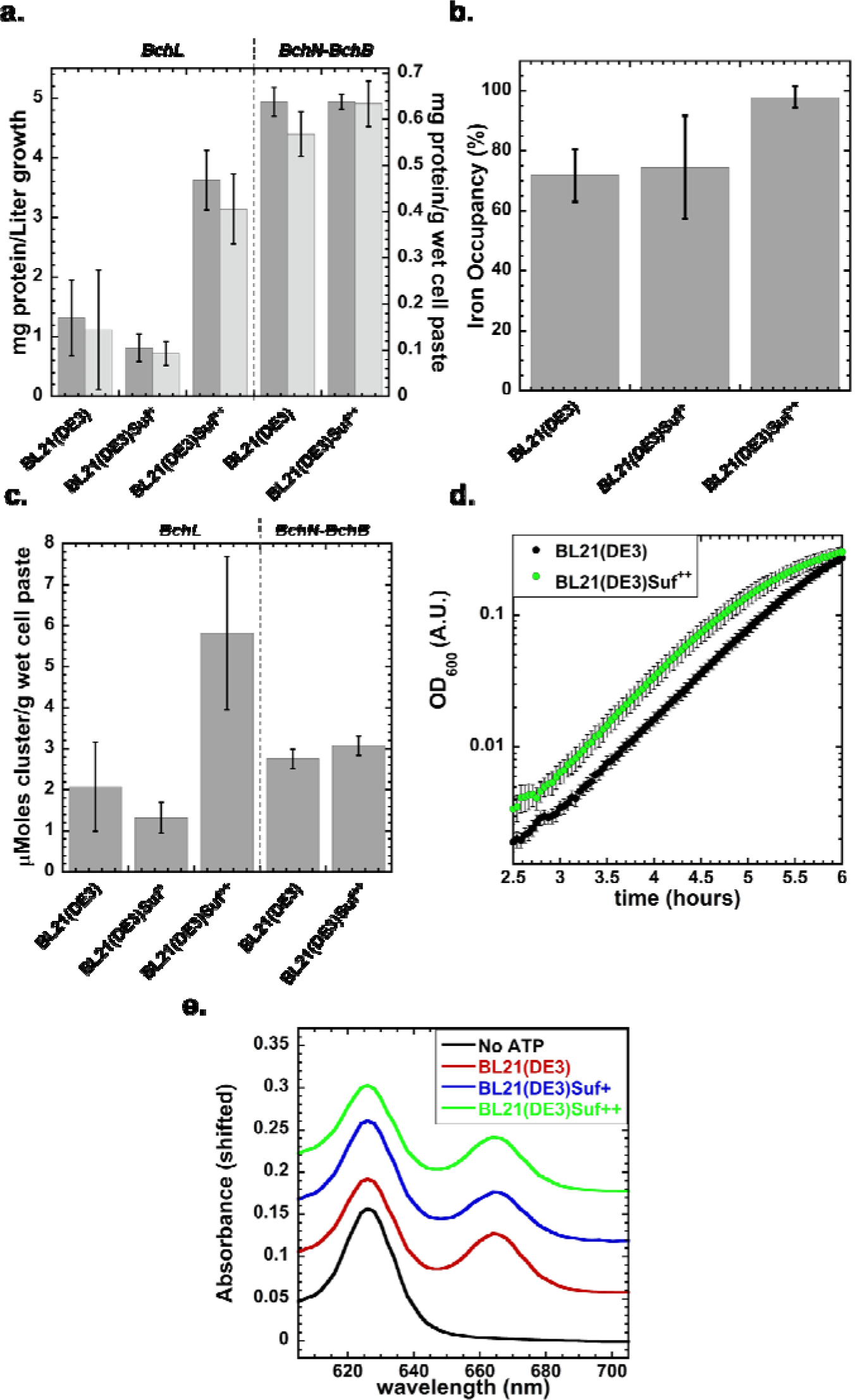
*SufFeScient* BL21(DE3) cells increase yield, [Fe-S] cluster formation/load ratio, exit lag phase earlier, and maintain enzyme activity. a) Quantification of the final yield of BchL and BchN-BchB proteins are plotted and show enhanced yield of the BchL protein in the BL21(DE3)Suf^++^ cells. No difference in the yield of BchN-BchB protein complex is observed. Left axis refers to dark gray bars, right axis refers to light gray bars. b) The total iron content in the purified BchL protein was measured and plotted as a percent of total occupancy. c) The ratio of [4Fe-4S] cluster per gram of wet cell paste is plotted for BchL and BchNBchB. d) Log-phase of Growth curves of BL21(DE3) (black circles) and BL21(DE3)Suf^++^ cells (green circles) that carry the BchL overexpression plasmid are plotted. Cells show similar doubling times (0.43 ± 0.10, and 0.41 ± 0.058 hours for BL21(DE3) cells and BL21(DE3)Suf^++^ cells, respectively, P= 0.61 from 2 tailed students t-test assuming unequal variance, n=9) but reach OD_600_=0.2 at different times ((5.91 ± 0.12, and 5.19 ± 0.17 hrs, for BL21(DE3) and BL21(DE3)Suf^++^ cells respectively, P=4.92E-08 from 2 tailed students t-test assuming unequal variance, n=9). e) BchL and BchN-BchB proteins purified from all three cell lines reduce Pchlide to Chlide. The lack of reduction in the absence of ATP is shown as control. The y-axis displays absorbance corresponding to the no-ATP dataset. The other three experimental traces have been numerically shifted up to better overlay the traces. Data in all panels show averaged results from three independent otherwise noted) is plotted.

We next tested whether the strain modifications influenced the overexpression or yield of the larger BchN-BchB hetero-tetramer, containing two [4Fe-4S] clusters. Unlike for BchL, BL21(DE3) and BL21(DE3) Suf^++^ overexpressed BchN-BchB to similar levels (Fig. 2e, f). Consequently, we did not obtain an enhancement in the yield of the BchN-BchB complex with BL21(DE3)Suf^++^ (Fig. 2i, 4.94±0.24 and 4.94±0.13 mg/L BchN-BchB from BL21(DE3), and BL21(DE3)Suf^++^, respectively).

We also compared the total iron occupancy within the purified BchL proteins from the three strains. Iron occupancy is shown here as (Molar [Fe] from purified protein determined by Dipyridyl colorimetric assay)/(theoretical Molar [Fe] assuming 4 Fe molecules per BchL dimer) X 100. Approximately 70–80% iron occupancy for the BchL dimer is observed when purified from BL21(DE3) and BL21(DE3)Suf^+^, compared to 100% occupancy when isolated from BL21(DE3)Suf^++^ (Fig. 3b). These data suggest that in addition to enhancing overall expression and increasing protein yields, the amount of protein carrying an intact [4Fe-4S] cluster is also higher in the BL21(DE3)Suf^++^ cells. BchN-BchB showed full occupancy when purified from any of the three strains (data not shown).

To explain why a ~3-fold increase in BchL production was observed in the BL21(DE3)Suf^++^ cells, we compared the [Fe-S] cluster formation/load ratio, which we define as the moles of [Fe-S] cluster recovered from purified enzyme per gram of wet cell mass (Fig. 3c). In this case, BL21(DE3)Suf^++^ cells generate and incorporate roughly three fold more (5 μmoles) clusters/g of wet cell mass compared to the other strains. This ratio does not scale for the BchN-BchB protein complex; the structural complexity of BchN-BchB and the mechanism of [Fe-S] cluster incorporation might be limiting protein folding, possibly explaining the lack of increased yields for the BchN-BchB complex. Interestingly, even before induction, there are consistent differences in the time required to exit lag phase between BL21(DE3) and BL21(DE3)Suf^++^, with the latter reaching optical densities appropriate for typical induction nearly one hour earlier than commercially available BL21(DE3), though their doubling times are not statistically different during log phase. (Fig. 3d). This may be due to increased overall cell health/viability even before induction of heterologous expression.

To ensure that the protein complexes purified from all the strains were active, we compared substrate reduction by their respective BchL and BchN-BchB proteins in vitro. Protochlorophyllide (substrate) and Chlide (reduced product) have unique spectral characteristics that are monitored through absorbance changes(7). Protochlorophyllide was monitored at its characteristic absorbance peak at 625 nm and formation of Chlide was captured at its absorbance peak at 668 nm (Fig. 3e). In the absence of ATP no formation of Chlide is observed (black trace, Fig. 3c), and addition of ATP triggers formation of a Chlide peak (red trace, Fig. 3c). The kinetics of substrate reduction are similar between the preparations when purified protein concentrations are normalized for reactions (0.24±.02, 0.27±.03, and 0.24±0.001 μM min^−1^ for proteins produced from BL21(DE3), BL21(DE3)Suf^+^, and BL21(DE3)Suf^++^ cells, respectively).

## DISCUSSION

Restoring the Suf [Fe-S] cluster biogenesis pathway and removing its transcriptional repression leads to a marked enhancement in the overexpression and purification of a [4Fe-4S] cluster protein. We call this BL21(DE3)Suf^++^ strain ‘SufFeScient’ for its potential application in improving overexpression and yields of other [Fe-S] cluster-containing proteins. The SufFeScient cells can improve yields in at least the case shown here; however, its utility must be experimentally determined for given target proteins since we do not have a sophisticated understanding of how subunit complexity, protein structure and local cluster environment impact biogenesis of target enzymes. For the BchL homodimer, the [4Fe-4S] cluster is exposed (Fig. 2a) whereas the two [4Fe-4S] clusters in the BchN-BchB complex are buried deep within the dimer interface in the context of a heterotetramer (Fig. 2a). Additionally, the relative efficiencies of the Isc versus Suf systems in incorporating [Fe-S] clusters into specific proteins are also not resolved but might account for the differences we observe.

BL21(DE3) is considered a workhorse strain for protein production because of the efficient control provided by T7 RNA polymerase integrated into its genome and the wide range of plasmid vectors containing T7 promoters used for protein overexpression or other biotechnological applications(12). The in-depth analysis of its genome sequence in 2009 provided a very important history of the progenitors of this strain as well as revealing several large deletions presumed to have been caused by UV irradiation that are specific to the BL21 lineage [see Table 4 of ref (31)]. Transcriptomics and metabolic modeling have provided additional insights into differences in the metabolic and transcriptional networks of this strain compared to other *E. coli*(11, 12, 17). Despite this wealth of knowledge, the genotype established in 2009 has not propagated to those listed by source companies and as a consequence the genotype listed in publications is typically incomplete [e.g. *E. coli* B F^−^*fhuA*2 *dcm ompT hsdS*(r_B_^−^ m_B_^−^) *gal* (λ DE3)]. Thus, the function of genes of interest in BL21(DE3) should be verified depending on experimental needs.

In fact, a previous study(28) noted the limitation of BL21(DE3) in producing other metal containing anaerobic respiratory enzymes. In this case the deficiency in producing some of these enzymes could be tracked to a nonsense mutation of the anaerobic transcription factor FNR and inefficient expression of proteins required for nickel transport. In addition, poor activity of some enzymes was also caused by a large 17, 247 bp deletion that removed the high affinity molybdate transport system, compromising the ability of BL21(DE3) to make the molybdenum cofactor necessary for function of several anaerobic respiratory enzymes. Of note, defects in activity for formate dehydrogenases-N and H were still detected even when these other systems were restored suggesting additional components in metalloenzyme synthesis are limiting in BL21(DE3). However, it seems unlikely that Suf dependent [Fe-S] cluster assembly was the step that was impaired since the Suf pathway is expressed at lower levels under anaerobic conditions(14).

In summary BL21(DE3) is a highly utilized host strain for over expressing [Fe-S] proteins and our correction of the deletion within the *suf* operon coupled with elevating its expression should provide researchers with another option for production of [Fe-S] proteins. As our knowledge for [Fe-S] cluster biogenesis pathway preferences for target proteins continues to expand, we will continue to optimize our ‘SufFeScient’ strain of BL21 to further improve [Fe-S] protein yields.

## MATERIALS AND METHODS

### Reagents and Buffers

Chemicals were purchased from Sigma-Millipore Inc. (St. Louis, MO), Research Products International Inc. (Mount Prospect, IL) and Gold Biotechnology Inc. (St. Louis, MO). Oligonucleotides were purchased from Integrated DNA Technologies (Coralville, IA). Enzymes for molecular biology were purchased from New England Biolabs (Ipswich, MA). All reagents and buffers used for protein purification were thoroughly degassed using alternating cycles of vacuum and nitrogen pressure on a home built Schlenk line apparatus. Anaerobic conditions were maintained via airtight syringes, excess reductant and a vinyl glove box (Coy Laboratories, MI) under a nitrogen (95%), hydrogen (5%) mix atmosphere.

### Strain construction

The genotypes of strains used in this study are shown in Table 1. A BL21(DE3) strain derivative (PK11466) was constructed to produce a functional *sufABCDSE* operon, but one that also lacks transcriptional repression by Fur. This was accomplished using P1 vir transduction to move a *cat*-P_*sufA*_(^−26^ATA^−24^ changed to ^−26^TAT^−24^) allele, which contains a mutation within the Fur binding site, from strain PK10882 to BL21(DE3) and selecting for growth on TYE agar plates containing 10 μg/ml chloramphenicol and 10 mM citrate. After streak isolating colonies twice on the same medium, colony PCR and DNA sequencing was carried out to confirm the genotype. Using the same method, BL21(DE3) strain derivative PK11465 was constructed to produce a functional *sufABCDSE* operon in which the Fur binding site within P_*sufA*_ is intact; the *cat*-P_sufA_ allele from PK10028 was moved to BL21(DE3) via P1 vir transduction.

Derivatives of MG1655 and BL21(DE3) were constructed to produce chromosomally-derived, epitope-tagged variants of SufB or SufS. First, a TAA stop codon was inserted directly after the 3XFLAG sequence on pIND4-3XFLAG(n) using Quikchange (Stratagene) to form pPK8629. Following PCR amplification of *cat* flanked by FLP recognition target (FRT) sites from pKD32 using primers containing BamHI and NdeI restriction sites, the PCR fragment was cloned into the same sites of pPK8629 to make pPK8630. To recombine 3XFLAG-TAA-FRT-*cat*-FRT directly before the native SufB (SufA/B in the case of BL21(DE3) or SufS stop codon, this construct was PCR amplified with primers containing homology to either region of the chromosome, electroporated into derivatives of MG1655 and BL21(DE3) that harbored pKD46, and selected for Cm^R^ (forming strains PK13230, PK13231, PK13232, and PK13233). After verification by DNA sequencing, *sufB*-3XFLAG-TAA-FRT-*cat*-FRT and *sufS*-3XFLAG-TAA-FRT-*cat*-FRT were respectively transduced via P1 vir from the MG1655 derivatives PK13230 and PK13231 to BL21(DE3), selecting for chloramphenicol resistance, to form strains PK13235 and PK13237.

**Table 1.**

| Strain or plasmid | Relevant genotype or phenotype | Reference or source |
| --- | --- | --- |
| **Strains** |  |  |
| BL21(DE3) | F<sup>-</sup> *hsdS gal* λDE3 | Laboratory stock |
| MG1655 | $\lambda$ F<sup>-</sup> *rph-1* | Laboratory stock |
| PK10882 | *zdh-3632::cat*, <sup>-26</sup>ATA<sup>-24</sup> bp relative to *sufA* TSS changed to <sup>-26</sup>TAT<sup>-24</sup> | (14) |
| PK11466 | BL21(DE3) but *zdh-3632::cat*, <sup>-26</sup>ATA<sup>-24</sup> bp relative to *sufA* TSS changed to <sup>-26</sup>TAT<sup>-24</sup> | This study |
| PK10028 | *zdh-3632::cat* | (14) |
| PK11465 | BL21(DE3) but *zdh-3632::cat* | This study |
| PK13230 | MG1655 but with *sufB*-3XFLAG-TAA-FRT-*cat*-FRT | This study |
| PK13231 | MG1655 but with *sufS*-3XFLAG-TAA-FRT-*cat*-FRT | This study |
| PK13232 | BL21(DE3) but with *sufA/B*-3XFLAG-TAA-FRT-*cat*-FRT | This study |
| PK13233 | BL21(DE3) but with *sufS*-3XFLAG-TAA-FRT-*cat*-FRT | This study |
| PK13235 | BL21(DE3) but with *sufB*-3XFLAG-TAA-FRT-*cat*-FRT derived from PK13230 | This study |
| PK13237 | BL21(DE3) but with *sufS*-3XFLAG-TAA-FRT-*cat*-FRT derived from PK13231 | This study |
| ZY-5 | *Rhodobacter sphaeroides* strain with deleted *bchL* used for Pchl<sub>a</sub> preparation | (7) |
| **Plasmids** |  |  |
| pIND4-3XFLAG(n) | *kan*, 3XFLAG | T. Donohue |
| pPK8629 | pIND4-3XFLAG(n) but with TAA inserted after 3XFLAG | This study |
| pPK8630 | pPK8629 but with FRT-*cat*-FRT cloned into BamHI and NdeI restriction sites | This study |
| pKD32 | FRT-*cat*-FRT | B. Wanner |
| pKD46 | Red recombinase expression plasmid | B. Wanner |
| BchL RSF-Duet1 | Plasmid used to overexpress BchL protein with a N-terminal 6x-Histidine tag and a TEV | This study |
|  | protease cleavage site |  |
| BchB RSF-Duet1 | Plasmid used to overexpress BchB protein with a N-terminal 6x-Histidine tag and a TEV protease cleavage site | This study |
| BchN pET-Duet1 | Plasmid used to overexpress BchN protein when coexpressed with BchB RSF-Duet1 | This study |
| NiCo21(DE3) | Commercial cell line | New England Biolabs |
| Lemo21(DE3) | Commercial cell line | New England Biolabs |
| C41(DE3) | Commercial cell line | Lucigen |
| Rosetta2(DE3)pLysS | Commercial cell line | EMD-Millipore-Novagen |
| BLR(DE3)pLysS | Commercial cell line | EMD-Millipore-Novagen |
| BL21(DE3)Ai | Commercial cell line | Thermo Fisher |
| BL21(DE3)codon plus | Commercial cell line | Agilent Technologies |

### Western blot analysis

Cultures were grown aerobically to an optical density at 600 nm of 0.1 in M9 minimal medium containing 0.2% glucose, 0.2% casamino acids, 1 mM MgSO_4_, 10 μg ml^−1^ ferric ammonium citrate, 4 μg ml^−1^ thiamine, and 0.1 mM CaCl_2_. Aliquots (1 ml) of cells were pelleted and levels of SufB-FLAG or SufS-FLAG were measured by Western blot analysis as previously described (16, 21) except that purified anti-DYKDDDDK epitope tag antibody (Biolegend) was used.

### Generation of protein synthesis plasmids

Plasmids used to overexpress BchL, BchN, and BchB were generated from PCR amplified *Rhodobacter sphaeroides* genomic DNA. BchL and BchB open reading frames were engineered to carry an N-terminal poly-histidine (6x-His) tag and 3C protease recognition sites and were cloned into pRSF-Duet1 using BamH1/Not1 and Sac1/Sal1 restriction sites, respectively. BchN contained no modifications and was cloned into a pET-Duet1 vector using Nde1/Kpn1 restriction sites.

### Generation of Pchlide

Pchlide was generated from a *Rhodobacter sphaeroides* ZY-5 strain harboring a deletion of the BchL gene (a kind gift from Dr. Carl Bauer, Indiana University).(23) ZY-5 cells were streaked from a glycerol stock onto agar plates containing RCV 2/3 medium, 15 g/L agar, and 0.5 mg/L kanamycin. Plates were grown overnight at 37°C and produced both white and brown colonies. A single brown colony was sub-streaked onto a RCV 2/3 agar plate to generate homogeneous brown colonies. Cells from the brown colonies were subsequently used to inoculate a 10 mL starter culture of RCV 2/3 medium with 0.5 mg/L kanamycin, grown overnight at 37°C, shaking at 200RPM. The entire 10 mL was then used to inoculate a fresh 125 mL culture with RCV 2/3 medium containing 0.05 mg/mL kanamycin. This culture was grown for 24 hours at 34°C, 130 rpm in a 250 mL flask completely covered in tin-foil to create a dark environment. Cells were centrifuged at 4,392 x g, and the green-colored spent medium was saved at 4°C until all growth steps were completed. The cell pellet was then resuspended in 500 mL fresh RCV 2/3 medium 0.05 mg/L kanamycin and grown for 24 hours at 34°C, 130 rpm in the dark. Cells were then pelleted and spent medium was saved at 4°C. 500mL fresh medium was added 2 additional times and grown for 24 hours each. Following growth, spent medium was centrifuged at 4,392 x g, 4°C for 1 hour to remove any remaining cells, and filtered through a white nylon 0.44 μm filter (Millipore, catalog #HNWP04700). Pchlide was extracted from the filtered spent medium with 1/3 volume di-ethyl ether in a standard fume hood. Extractions were performed in 500 mL increments in a 1L separatory funnel taking care to vent built up pressure from volatile ether after vigorous shaking. Organic and aqueous phases were allowed to separate for 15 minutes before collecting the green colored organic phase. The collected organic phase was then centrifuged at 4122 x g at 4°C for 5 minutes to further separate any lipid or aqueous contamination which appears as transparent droplets below the green ether layer. The ether layer was then decanted and evaporated to dryness under a stream of nitrogen in a standard fume hood. Water droplets formed on the inside and outside of the vessel were allowed to evaporate inside the fume hood. The resulting dark green (nearly black) flaky material was then resuspended in 250 μL DMSO/500 mL spent medium extraction. The suspended Pchlide was aliquoted and stored in opaque brown “Graduated Amber” 1.5 mL conical tubes (Fisherbrand; Catalog #05-408-134) and stored covered in tin foil at 4°C. Concentration of Pchlide was determined using 4 dilutions, three replicates each in 80 % acetone using the molar extinction coefficient 30,400 M^−1^cm^−1^.

### Protein synthesis and purification

Bacterial cells transformed with the respective plasmid for recombinant protein overexpression were grown in 2.8 L baffled-flasks containing 1 L autoclaved Luria Broth medium supplemented with appropriate antibiotic (100 mg/L ampicillin and/or 50 mg/L kanamycin), 1 mM ferric citrate and 1 mM L-cysteine. Large cultures were inoculated with a suspension of all the colonies from a fresh transformation, and cultures were then grown aerobically at 37°C shaking at 200 rpm. Protein synthesis was induced at OD600=0.6 with 35 μM IPTG from a freshly prepared 100 mM stock, and cultures were shifted to 25°C shaking at 150 RPM overnight (~17 hr). Cells were harvested via centrifugation (4200 rpm, 20 minutes, 4°C) after a 3-hour incubation with sodium dithionite (0.3 g/L growth) in 1 L airtight centrifuge tubes at 17°C with no shaking. All subsequent steps were performed in the glove box, or in airtight septum sealed bottles under positive nitrogen pressure unless otherwise noted. Cell pellets were resuspended in 25 mL/L growth of degassed STD buffer (100 mM HEPES pH 7.5, 150 mM NaCl, 1.7 mM sodium dithionite) after decanting spent medium inside of a glovebox. Resuspended cells were transferred to a septa-sealed glass bottle and stored at −20 °C until lysis.

Cell lysis was performed using lysozyme (0.5 mg/ml) for 30 minutes at room temperature in septum sealed bottles under positive nitrogen pressure. Cells were then sonicated inside a glovebox for 3 minutes on ice (Branson sonifier, 50 % duty cycle, 60 seconds on, 60 seconds off for 3 cycles). Cell lysates were clarified via centrifugation (37,157 x g, 60 minutes) in gasket-sealed 40 mL centrifuge tubes. Clarified lysates were loaded onto a Nickel-nitrilotriacetic acid (Ni^2+^-NTA) column (Thermo Scientific). Ni^2+^-NTA column volumes were 1.6 mL resuspended beads per liter growth. Ni^2+^-NTA columns were equilibrated with 10 column volumes (CVs) STD buffer using a peristaltic pump at 2.0 mL/min. After loading the clarified lysate at 2.0 ml/min, the column was then washed with 10 CVs STD buffer containing 20 mM imidazole. Proteins were eluted into a septum sealed bottle using 30 mL STD buffer containing 250 mM imidazole. Eluted BchN-BchB protein were concentrated using an Amicon 30 kDa molecular weight cut-off spin concentrator (Sigma-Millipore Inc., St. Louis, MO), centrifuged for 8 minutes at 4122 x g at 4°C. BchL protein eluted from the Ni^2+^-NTA column were subsequently loaded onto a 10 mL Q-Sepharose column (GE Healthcare) using an AKTA pure FPLC (GE Healthcare) after first washing with 10 CVs of 150 mM HEPES pH 7.5, 1 M NaCl (buffer B) followed by equilibration with 10 CVs of 150mM HEPES pH 7.5 at 2.0 mL/min. BchL protein eluate was subsequently loaded at 2.0 ml/min and non-specifically bound impurities were removed by washing the column with 10 CV STD buffer. BchL was next eluted with 140 ml linear gradient: 100% STD buffer (line A) to 100% buffer B (line B), collecting 1.8 mL fractions inside a glovebox. Fractions containing BchL protein were concentrated using a spin concentrator as described above for the BchN-BchB proteins. Proteins were aliquoted into 1.2mL cryo-tubes (Cat430487 Corning) in the glove box, which have a gasket-sealed cap, removed from the glove box and flash frozen using liquid nitrogen. The protein contained in cryo-tubes were stored under liquid nitrogen.

Comparison of protein induction and purification using SDS-PAGE analysis 10 % SDS polyacrylamide gels were used to visualize protein samples. Un-induced and induced samples (1.0 mL and 0.5 mL, respectively) were centrifuged in a table-top centrifuge (13,222 x g, 60 sec, 4°C), and the supernatant was decanted. Sample cell pellets were re-suspended in 100 μL water. Tenfold dilutions of resuspended cells were used to record the OD_600_ and this value was used to normalize all sample optical densities. Equivalent units of OD_600_ of undiluted samples were then diluted to 100μL and mixed with 100 μl of 2X SDS Laemmli dye and boiled for 10 min. For all the gels shown in this paper, 10 μl of boiled cell samples were analyzed on the SDS-PAGE gels. PageRuler Plus Prestained Ladder (Thermo Scientific) was used as a protein size ladder for reference.

### Assay for substrate reduction by DPOR

Reduction of Pchlide to Chlide was measured spectroscopically by mixing BchN-BchB protein (3 μM tetramer), BchL-protein (9 μM dimer), and 35 μM Pchlide, in the absence or presence of ATP (3 mM) in STD buffer + 10mM MgCl_2_. 40 μl of these reactions were quenched with 160 μl of 100 % acetone (80% v/v final concentration). The acetone extraction was then centrifuged in a table top centrifuge (13,226 x g for 4 minutes) to pellet precipitated protein components. 160 μl of the supernatant was transferred to a cyclic olefin half-area well plate (catalog #4680 Corning) and absorbance scans from 600 nm to 725 nm were recorded on a SpectraMax i3x plate reader (Molecular Devices). Chlide appearance was quantified using its molar extinction coefficient 74,900 M^−1^cm^−1^ at 666 nm.

### Protein and Iron-content determination

Protein concentrations were determined using the Bradford Assay reagent (BioRad) and Bovine Serum Albumin (GoldBio) as a standard. Iron content was determined by colorimetric assay using 2’2’dipyridal absorbance at 520nm and Fe(NH_4_)_2_(SO_4_)_2_ (Fisher) as a standard after sample denaturation and iron reduction (60 minute exposure to 5% HCl and boiling, followed by exposure to excess (10%) hydroxylamine.)

### Growth curve generation

BL21(DE3) and BL21(DE3)Suf^++^ cell lines were transformed with BchL plasmid and plated on LB-Agar containing 0.5mg/L (1X) kanamycin. 6 individual colonies from both cell lines were picked and used to inoculate a 5 mL overnight starter culture (LB + 0.5mg/L kanamycin). Starter cultures were normalized to OD600 =0.01 with LB + 0.5mg/L kanamycin. 2 uL of OD600 corrected starter cultures were used to inoculate 200 uL growths in a polystyrene 96-Well cell-culture plate (Cole-Palmer). 6 wells containing only 200 uL LB + 0.5mg/L kanamycin served as a blank and negative control. Cells were grown inside a SpectraMax i3x plate reader at 37°C, shaken for 10 seconds before each read. OD600 was measured every 2.5 minutes. Blank measurements were averaged, and individual growth curves were corrected by subtracting averaged blank values at each time point, then averaged after correction for Figure 3d. Doubling rates (“r”) were calculated from t=2 and t=4 hours using r=(ln[OD2/OD1])/(t2-t1). Doubling time was calculated using ln(2)/r.

## ACKNOWLEDGEMENTS

E.I.C., and E.L.M. performed experiments. E.A. and P.J.K. conceived the experiments. All authors contributed to writing the manuscript. This work was supported by a grant from the Department of Energy, Office of Basic Energy Sciences DE-SC0017866 to E.A. and NIH grant R01-GM115894 to P.J.K.

## REFERENCES

1. Beinert, H. 2000. Iron-sulfur proteins: ancient structures, still full of surprises. J Biol Inorg Chem 5: 2–15.

2. Blanc, B., C. Gerez, and S. Ollagnier de Choudens. 2015. Assembly of Fe/S proteins in bacterial systems: Biochemistry of the bacterial ISC system. Biochim Biophys Acta 1853: 1436–1447.

3. Boyd, E. S., K. M. Thomas, Y. Dai, J. M. Boyd, and F. W. Outten. 2014. Interplay between oxygen and Fe-S cluster biogenesis: insights from the Suf pathway. Biochemistry 53: 5834–5847.

4. Brocker, M. J., S. Schomburg, D. W. Heinz, D. Jahn, W. D. Schubert, and J. Moser. 2010. Crystal structure of the nitrogenase-like dark operative protochlorophyllide oxidoreductase catalytic complex (ChlN/ChlB)_2_. J Biol Chem 285: 27336–27345.

5. Daegelen, P., F. W. Studier, R. E. Lenski, S. Cure, and J. F. Kim. 2009. Tracing ancestors and relatives of *Escherichia coli* B, and the derivation of B strains REL606 and BL21(DE3). J Mol Biol 394: 634–643.

6. Fontecave, M. 2006. Iron-sulfur clusters: ever-expanding roles. Nat Chem Biol 2: 171–174.

7. Fujita, Y., and C. E. Bauer. 2000. Reconstitution of light-independent protochlorophyllide reductase from purified bchl and BchN-BchB subunits. In vitro confirmation of nitrogenase-like features of a bacteriochlorophyll biosynthesis enzyme. J Biol Chem 275: 23583–23588.

8. Grawert, T., J. Kaiser, F. Zepeck, R. Laupitz, S. Hecht, S. Amslinger, N. Schramek, E. Schleicher, S. Weber, M. Haslbeck, J. Buchner, C. Rieder, D. Arigoni, A. Bacher, W. Eisenreich, and F. Rohdich. 2004. IspH protein of Escherichia coli: studies on iron-sulfur cluster implementation and catalysis. J Am Chem Soc 126: 12847–12855.

9. Jeong, H., V. Barbe, C. H. Lee, D. Vallenet, D. S. Yu, S. H. Choi, A. Couloux, S. W. Lee, S. H. Yoon, L. Cattolico, C. G. Hur, H. S. Park, B. Segurens, S. C. Kim, T. K. Oh, R. E. Lenski, F. W. Studier, P. Daegelen, and J. F. Kim. 2009. Genome sequences of *Escherichia coli* B strains REL606 and BL21(DE3). J Mol Biol 394: 644–652.

10. Kiley, P. J., and H. Beinert. 2003. The role of Fe-S proteins in sensing and regulation in bacteria. Curr Opin Microbiol 6: 181–185.

11. Kim, H., S. Kim, and S. H. Yoon. 2018. Metabolic network reconstruction and phenome analysis of the industrial microbe, *Escherichia coli* BL21(DE3). PLoS One 13: e0204375.

12. Kim, S., H. Jeong, E. Y. Kim, J. F. Kim, S. Y. Lee, and S. H. Yoon. 2017. Genomic and transcriptomic landscape of *Escherichia coli BL21*(DE3). Nucleic Acids Res 45: 5285–5293.

13. Kriek, M., L. Peters, Y. Takahashi, and P. L. Roach. 2003. Effect of iron-sulfur cluster assembly proteins on the expression of *Escherichia coli* lipoic acid synthase. Protein Expr Purif 28: 241–245.

14. Mettert, E. L., and P. J. Kiley. 2014. Coordinate regulation of the Suf and Isc Fe-S cluster biogenesis pathways by IscR is essential for viability of *Escherichia coli*. J. Bacteriol. 196: 4315–4323.

15. Mettert, E. L., and P. J. Kiley. 2015. How Is Fe-S Cluster Formation Regulated? Annu Rev Microbiol 69: 505–526.

16. Mettert, E. L., F. W. Outten, B. Wanta, and P. J. Kiley. 2008. The Impact of O_2_ on the Fe-S Cluster Biogenesis Requirements of *Escherichia coli* FNR. Journal of Molecular Biology 384: 798–811.

17. Monk, J. M., A. Koza, M. A. Campodonico, D. Machado, J. M. Seoane, B. O. Palsson, M. J. Herrgard, and A. M. Feist. 2016. Multi-omics Quantification of Species Variation of *Escherichia coli* Links Molecular Features with Strain Phenotypes. Cell Syst 3: 238–251 e212.

18. Moser, J., C. Lange, J. Krausze, J. Rebelein, W. D. Schubert, M. W. Ribbe, D. W. Heinz, and D. Jahn. 2013. Structure of ADP-aluminium fluoride-stabilized protochlorophyllide oxidoreductase complex. Proc Natl Acad Sci U S A 110: 2094–2098.

19. Muraki, N., J. Nomata, K. Ebata, T. Mizoguchi, T. Shiba, H. Tamiaki, G. Kurisu, and Y. Fujita. 2010. X-ray crystal structure of the light-independent protochlorophyllide reductase. Nature 465: 110–114.

20. Nakamura, M., K. Saeki, and Y. Takahashi. 1999. Hyperproduction of recombinant ferredoxins in *Escherichia coli* by coexpression of the *ORF1-ORF2-iscS-iscU-iscA-hscB-hs cA-fdx-ORF3* gene cluster. J Biochem 126: 10–18.

21. Nesbit, A. D., J. L. Giel, J. C. Rose, and P. J. Kiley. 2009. Sequence-specific binding to a subset of IscR-regulated promoters does not require IscR Fe-S cluster ligation. Journal of Molecular Biology 387: 28–41.

22. Nomata, J., M. Kitashima, K. Inoue, and Y. Fujita. 2006. Nitrogenase Fe protein-like Fe-S cluster is conserved in L-protein (BchL) of dark-operative protochlorophyllide reductase from *Rhodobacter capsulatus*. FEBS Lett 580: 6151–6154.

23. Nomata, J., T. Kondo, T. Tamiaki, S. Itoh, and Y. Fujita. 2014. Dark-operative protochlorophyllide oxidoreductase generates substrate radicals by an iron-sulphur cluster in bacteriochlorophyll biosynthesis. Sci Rep 4. 5455.

24. Nomata, J., T. Ogawa, M. Kitashima, K. Inoue, and Y. Fujita. 2008. NB-protein (BchN-BchB) of dark-operative protochlorophyllide reductase is the catalytic component containing oxygen-tolerant Fe-S clusters. FEBS Lett 582: 1346–1350.

25. Nomata, J., L. R. Swem, C. E. Bauer, and Y. Fujita. 2005. Overexpression and characterization of dark-operative protochlorophyllide reductase from *Rhodobacter capsulatus*. Biochim Biophys Acta 1708: 229–237.

26. Outten, F. W. 2015. Recent advances in the Suf Fe-S cluster biogenesis pathway: Beyond the Proteobacteria. Biochim Biophys Acta 1853: 1464–1469.

27. Outten, F. W., O. Djaman, and G. Storz. 2004. A *suf* operon requirement for Fe-S cluster assembly during iron starvation in *Escherichia coli*. Mol Microbiol 52: 861–872.

28. Pinske, C., M. Bonn, S. Kruger, U. Lindenstrauss, and R. G. Sawers. 2011. Metabolic deficiences revealed in the biotechnologically important model bacterium *Escherichia coli* BL21(DE3). PLoS One 6: e22830.

29. Py, B., and F. Barras. 2010. Building Fe-S proteins: bacterial strategies. Nat Rev Microbiol 8: 436–446.

30. Roche, B., L. Aussel, B. Ezraty, P. Mandin, B. Py, and F. Barras. 2013. Iron/sulfur proteins biogenesis in prokaryotes: formation, regulation and diversity. Biochim Biophys Acta 1827: 455–469.

31. Studier, F. W., P. Daegelen, R. E. Lenski, S. Maslov, and J. F. Kim. 2009. Understanding the differences between genome sequences of *Escherichia coli* B strains REL606 and BL21(DE3) and comparison of the *E. coli* B and K-12 genomes. J Mol Biol 394: 653–680.

32. Takahashi, Y., and U. Tokumoto. 2002. A third bacterial system for the assembly of iron-sulfur clusters with homologs in archaea and plastids. J Biol Chem 277: 28380–28383.

33. Tokumoto, U., S. Kitamura, K. Fukuyama, and Y. Takahashi. 2004. Interchangeability and distinct properties of bacterial Fe-S cluster assembly systems: functional replacement of the isc and suf operons in *Escherichia coli* with the nifSU-like operon from *Helicobacter pylori*. J Biochem 136: 199–209.

34. Tsai, C. L., and J. A. Tainer. 2018. Robust Production, Crystallization, Structure Determination, and Analysis of [Fe-S] Proteins: Uncovering Control of Electron Shuttling and Gating in the Respiratory Metabolism of Molybdopterin Guanine Dinucleotide Enzymes. Method Enzymol 599: 157–196.

